# Prediction of the effect of pH on the aggregation and conditional folding of intrinsically disordered proteins with SolupHred and DispHred

**DOI:** 10.1101/2021.06.08.447585

**Authors:** Valentín Iglesias, Carlos Pintado-Grima, Jaime Santos, Marc Fornt, Salvador Ventura

## Abstract

Proteins microenvironments modulate their structures. Binding partners, organic molecules, or dissolved ions can alter the protein’s compaction, inducing aggregation or order-disorder conformational transitions. Surprisingly, bioinformatic platforms often disregard the protein context in their modeling. In recent work, we proposed that modeling how pH affects protein net charge and hydrophobicity might allow us to forecast pH-dependent aggregation and conditional disorder in intrinsically disordered proteins (IDPs). As these approaches showed remarkable success in recapitulating the available bibliographical data, we made these prediction methods available for the scientific community as two user-friendly web servers. SolupHred is the first dedicated software to predict pH-dependent aggregation, and DispHred is the first pH-dependent predictor of protein disorder. Here we dissect the features of these two software applications to train and assist scientists in studying pH-dependent conformational changes in IDPs.

## 1 Introduction

Most proteins reside in crowded environments inside the cells and establish different interactions with various partners: other proteins, complex carbohydrates, nucleic acids, inorganic ions, and other metabolites. Intrinsically disordered proteins (IDPs) are particular polypeptides that do not present a defined stable, compact structure. Instead, they might transiently populate different degrees of conformations, ranging from fully structured to completely disordered, with a continuum of partially structured folds in between [1]. Their disordered nature is maintained by a compositional bias favoring charged, polar, and proline residues and disfavouring aliphatic amino acids [2]. In essence, by avoiding the hydrophobic effect that drives protein folding and aggregation, they preserve their disordered nature and structural plasticity.

Computational approaches have co-evolved with our understanding of IDPs’ biology, becoming powerful platforms to rationalize, model, and identify their conformational determinants and effectors. This has resulted in a versatile toolbox of computational software that allows the rapid and cost-effective analysis of disordered sequences, even to the point of engineering new protein variants *à la carte* [3–5]. However, IDP’s low cooperativity and high solvent exposition make them highly sensitive to environmental fluctuations [6], influencing their fundamental properties and compromising *in silico* predictions. Nevertheless, this influence is often disregarded or barely parametrized, and most bioinformatic approaches perform their analysis assuming standard conditions: physiological temperature, pressure, salt concentration, and neutral pH. However, these conditions do not apply for multiple proteins such as those residing in acidic or basic compartments (i.e., lysosomes, storage granules, gastrointestinal tract, pancreas) [7–9] or in organisms adapted to live in extreme conditions (halophiles, thermophiles or cryophiles, alkaliphiles or acidophiles...) [10–12]. Besides, the production and formulation of protein-based drugs often involve drastic changes in the solution conditions. This is especially true for the pH, since purification protocols often require distinct buffers, which might differ significantly in their pHs. Furthermore, most protein biotherapeutics are purified, stored, or formulated at non-neutral pH, as in antibodies, with over 65% of them being formulated at acidic pH [13,14]. Thus, it appeared necessary to integrate pH into *in silico* calculations to anticipate the experimental behavior of IDPs. In previous works, we rationalized the influence of pH on IDPs’ disordered states and aggregation propensities and modeled their dependence on this parameter in such reactions [15–17]. Our models succeeded in identifying pH-dependent aggregation and order-disorder transitions, as reported in the literature. Then, we made these approaches available to the scientific community as easy-to-use web servers, SolupHred and DispHred, providing rapid and publically accessible methods to evaluate the adequate pH intervals for their particular IDPs and applications of interest. In this chapter, we describe the methodological pipeline behind SolupHred and DispHred and illustrate their use.

## 2 Materials

It is well established that pH can vary the charge of ionizable amino acids through protonation or deprotonation. However, its influence on hydrophobicity is generally disregarded, even if hydrophobicity is a primary determinant of protein folding and aggregation processes. Accordingly, we reasoned that the effect of pH on IDPs compaction could be modeled by accounting for these two pH-dependent physicochemical properties: hydrophobicity and net charge. By including the pH-dependency of hydrophobicity, we increased our predictive potential compared to current methodologies that only account for the impact of pH on the protein charge. To do so, we exploited a recent study where Zamora and coworkers [18] developed a pH-dependent amino acid lipophilicity scale by computing the *n*-octanol/water partition for amino acids in protein-like environments using continuum solvation calculations. Even if lipophilicity and hydrophobicity are not strictly equivalent terms, the aforementioned scale showed a good correlation with hydrophobicity-based scales widely used in applications of disorder prediction [17,19–21] or aggregation [15,22]. Therefore, it will serve as a legit proxy for the calculus of protein hydrophobicity. In parallel, the net charge of individual amino acids at different pHs was calculated using the Henderson-Hasselbalch equation.

### 2.1 Modeling pH-dependent aggregation of intrinsically disordered proteins with SolupHred

#### 2.1.1 Algorithm rationale

IDPs constituted a compelling testing dataset as they lack any structural element that could dim the impact of the physicochemical properties of the primary sequence we aimed to study. In a previous study, S. Brocca’s lab engineered three protein variants of a model IDP by reversing the sign of charged residues already present in the wild-type sequence [23]. This rendered proteins with different net charges and isoelectric points (pI), for which they experimentally characterized their solubility at different pHs. In collaboration with Brocca’s lab, we calculated lipophilicity and net charge for each experimentally derived datapoint and plotted the dispersion into a 3D plot, which delineated the flat 2D surface of a plane in a Euclidean space. Accordingly, the data were fitted into a bivariate polynomial model with a quadratic component (Equation 1) through the non-linear least-squares approach of the Scipy Python module [24].

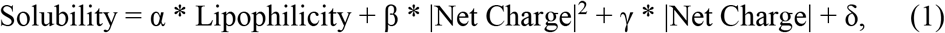

The experimentally observed values highly correlated with the model predictions (p-value < 0.00001), whereas considering charge as the only pH-dependent factor ultimately failed to provide an accurate description of the data.

#### 2.1.2 Pipeline

SolupHred calculates lipophilicity and net charge for the input sequences at the requested pHs, as described previously, and applies Equation 1 to retrieve the maximum and minimum solubilities at defined pHs (Note 1). The server requires introducing or uploading a valid FASTA formatted sequence file, along with the pH interval to analyze. For each pH datapoint, the server uses a size-dependent sliding window that accounts for the influence of neighboring residues, as previously described for widely-employed sequence-based prediction methods [22] (Note 2). Mean lipophilicity of the sequence is then calculated as the average of all local lipophilicities. Global net charge is calculated using the Henderson-Hasselbalch equation for individual residues and summing the individual components. For every pH, SolupHred combines the obtained mean lipophilicity and net charge using Equation 1. The results are retrieved as means of two user-friendly tables, the first one specifying the main results and the second one with the raw data, an intelligible figure with the pHs for predicted top and least 10% solubility (Note 3) and a machine-and-human-readable array-based JavaScript Object Notation (JSON) object with the complete data (Figure 1). Alternatively, users can select to predict solubility at a single pH, where a simplified output will be displayed.

**Figure 1.**
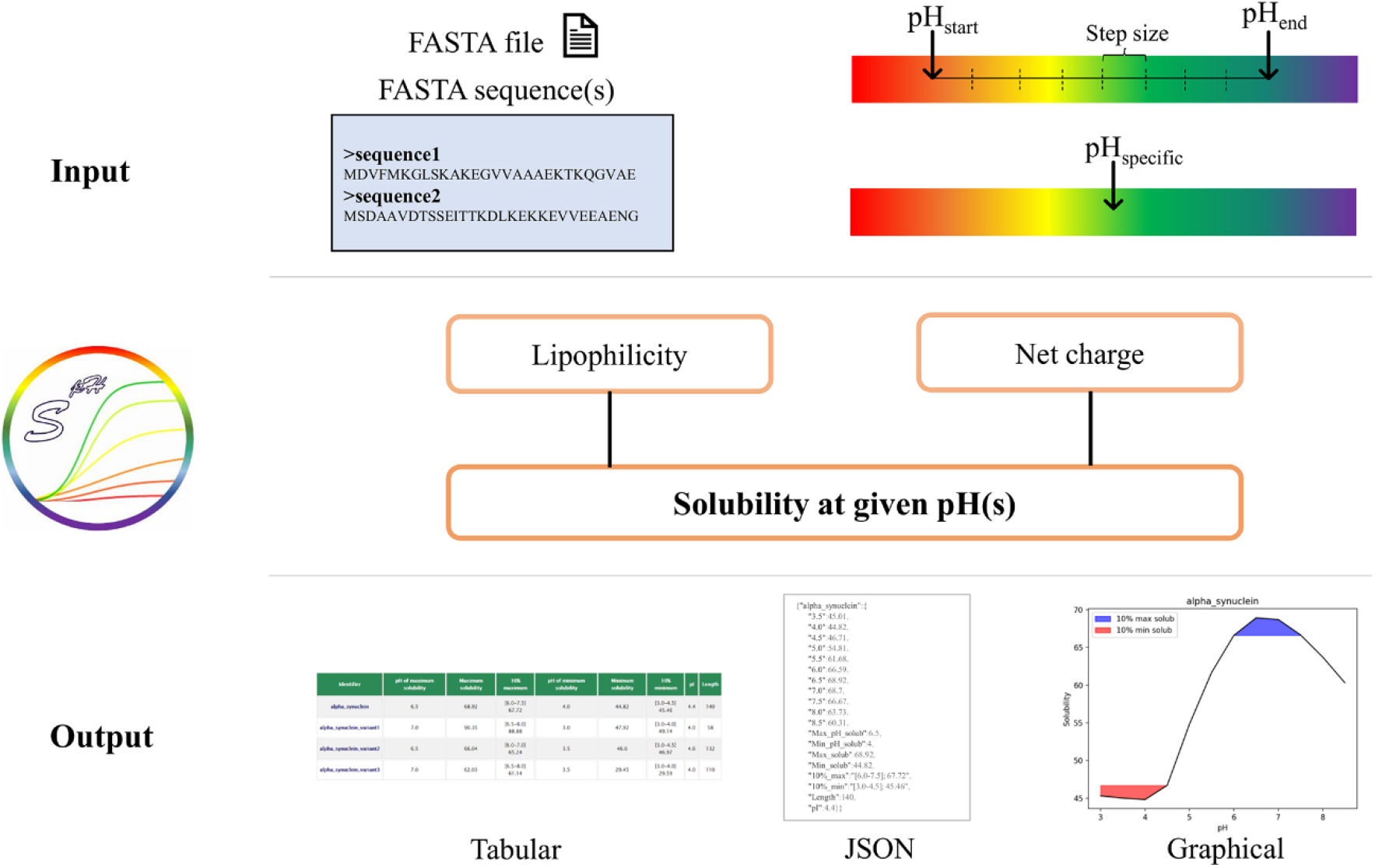
SolupHred pipeline: SolupHred acquires input sequences and a pH or pH range for analysis. SolupHred will calculate the lipophilicity and net charge for the specified options and apply Equation 1 to determine its solubility. SolupHred will retrieve the output as easy-to-interpret tables, graphs, and a machine-readable JSON file.

SolupHred web server interface is built using a combination of HTML, CSS and Javascript, processed by a Django CGI and a backend script written in Python that uses Python 3.7 as the interpreter. SolupHred uses Python numpy, scipy and matplotlib data-analysis libraries to process the data [25,24].

### 2.2 Predicting pH-dependent order-disorder transitions in intrinsically disordered proteins with DispHred

#### 2.2.1 Algorithm rationale

In 2000, Uversky and coworkers described that IDPs populated a specific region in the charge-hydropathy (C-H) space diagram [2], and an empirically defined boundary line was able to discriminate them from folded proteins. This rationale has since been applied in state-of-the-art disorder prediction algorithms [26]. Since both hydrophobicity and net charge are pH-dependent, we applied the same rationale used in SolupHred to extend the C-H predictive potential to the entire pH range. We collected all available bibliographical data on pH-dependent order-disorder transitions in IDPs and plotted their pH, calculated lipophilicity (as a proxy for hydrophobicity), and net charge into a 3D chart. Our pH-dependent approach was able to discriminate between ordered and disordered sequences. To achieve the best possible order-disorder boundary condition, we applied a support vector machine (SVM) learning approximation (Equation 2) [27,28]. SVM allows a certain degree of misclassified data points without overfitting, achieving the maximal separation between the two states and providing a margin that is employed as a confidence interval.

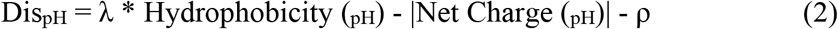

The experimentally obtained data could be predicted with great accuracy, whereas, again, considering only net charge as a pH-dependent was unable to foresee the degree of disorder in each particular pH condition.

#### 2.2.2 Pipeline

DispHred utilizes the calculation of pH-dependent hydrophobicity and net charge to extend the C-H space-analysis to different pH ranges. This allows it to discriminate between ordered and disordered states of proteins in any given condition. The server requires introducing a valid FASTA formatted sequence or a valid Uniprot Accession number [29], the pH interval to analyze, and pre-fixed window size (Note 4). For each requested pH, DispHred calculates hydrophobicity as the average of all windows lipophilicities and the net charge per residue by dividing the overall charge by the protein length. Dis_pH_ score is then obtained by contrasting the hydrophobicity and net charge with the boundary condition defined in Equation 2. Sequences with positive Dis_pH_ scores are predicted as ordered, while those with negative are predicted to be disordered at that pH. The server has a margin of ± 0.02, which is used as a confidence interval, therefore predictions that at a specific pH score 0 ± 0.02 should not be regarded as ambiguous. DispHred retrieves calculations easily through a table with the raw scores and a graph plotting the Dis_pH_ score variation across the requested pH range. The server provides the total output in i) a table and ii) a graph depicting disorder-prediction against pH, iii) a resume of the pH-dependent predictions along the sequence for the specified pH range. Alternatively, users may select to predict the disordered state at a defined pH. In that case, a modified output will be displayed, showing i) a table with the calculations for Dis_pH_ score, hydrophobicity, and net charge accompanied by ii) a plot that displays Dis_pH_ score per residue along the sequence and iii) a table with the windows scores for the sequence profile. Additionally, users can obtain the complete calculations as a JSON object or download the complete dataset in a ZIP file. A scheme of DispHred’s pipeline is shown in Figure 2.

**Figure 2.**
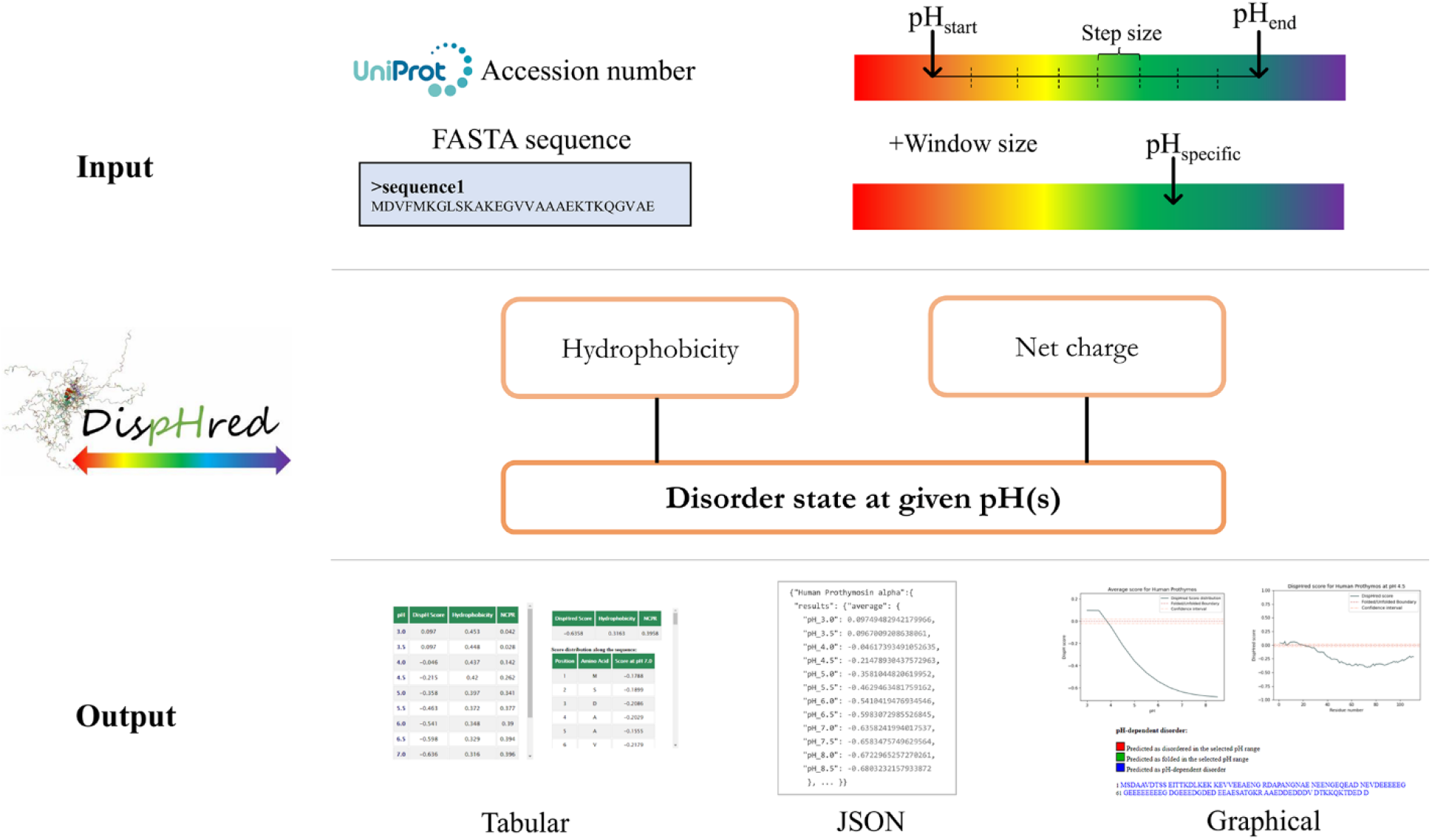
DispHred pipeline: DispHred takes Uniprot Accession number or a FASTA sequence as an input along with the specified pH or pH range for the calculations. On the backend, it will calculate the hydrophobicity and net charge for each pH and apply Equation 2 to perform the prediction. The results are presented in a user-friendly manner as a table and graphs that summarize the main results and an easy to parse JSON file.

The DispHred web server interface is built using HTML, CSS, and Javascript technologies; a Django CGI processes the input and outputs, and the backend script is written in Python uses Python 3.7 as the interpreter. In addition, DispHred uses widely used Python data-analysis libraries numpy, scipy, and matplotlib to process the data [25,24].

## 3 Methods

Both web servers are designed as easy-to-use approaches to support the study of pH-dependent protein folding and aggregation and as handy tools to explore the response of IDPs at different pHs or the assistance of engineering pH-responsive proteins. Therefore, they are freely available for academic purposes and do not require previous login or registration. SolupHred can be accessed at https://ppmclab.pythonanywhere.com/SolupHred and DispHred at https://ppmclab.pythonanywhere.com/DispHred.

### 3.1 SolupHred web server

#### 3.1.1 Frontpage and running options

SolupHred web server displays a straightforward and user-friendly interface (Figure 3). Four different sections arise from the top in a navigation bar. Users may access a brief explanation of the method by clicking *Help* or to the two main publications by clicking *References.* For any further inquiries, SolupHred provides a *Contact* section with a correspondence email address. The main page places users in the Submission subsection, where all input parameters and data are introduced and defined. SolupHred allows users to paste or upload the query sequences, provided they are FASTA-formatted (Notes 5 and 6).

**Figure 3.**
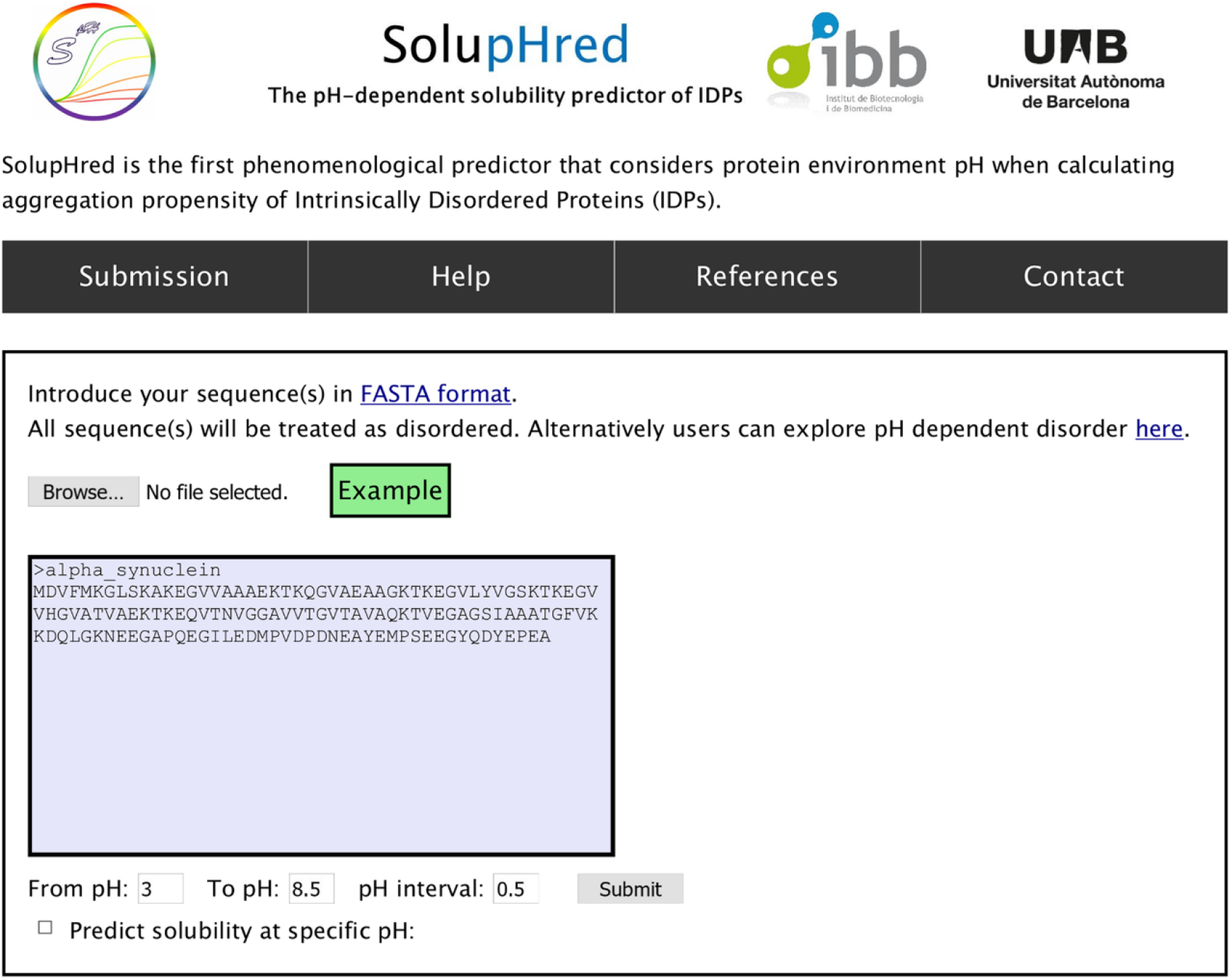
SolupHred front page: On top of the page, SolupHred displays four headers: *Submission, Help, References,* and *Contact.* Under *Submission,* users can paste or upload their FASTA-formatted sequence(s) and define a pH or pH range to perform the analysis. On clicking the *Example* button, the server will pre-populate the human α-synuclein sequence and a pH range for which its solubility has been shown to change significantly.

A pH range for the analysis must be selected by filling the *“From pH”* and *“To pH”* boxes, and a step size *pH interval* should be defined (Notes 7 and 8). Users can try the server by pressing the “Example” link, which will pre-populate the window with the sequence of α-synuclein and a pH-range for which its solubility has been shown to diverge [30,31]. Alternatively, users can select to predict the solubility of proteins at a specific pH by clicking the checkbox next to *“Predict solubility at a specific pH”.* Pressing *Submit* will compute the input options and start the solubility analysis.

#### 3.1.2 Output and evaluation

After SolupHred completes the task, it redirects users to an output page. The main output for a single protein, and more than one pH, displays two tables and a plot depicting solubility against pH (Figure 4). The central table summarizes at a glance the most relevant results: the pH at which the protein is predicted to be most and least soluble (Note 3), along with the scores, the protein length, and the calculated isoelectric point. Immediately below, the screen is divided into two parts. The left side depicts a plot representing the predicted solubility variation along with the specified pH range, with the 10% maximum solubility colored in blue and the 10% minimum in red. A table shows the numerical value for the prediction at every requested pH on the right side of the screen.

**Figure 4.**
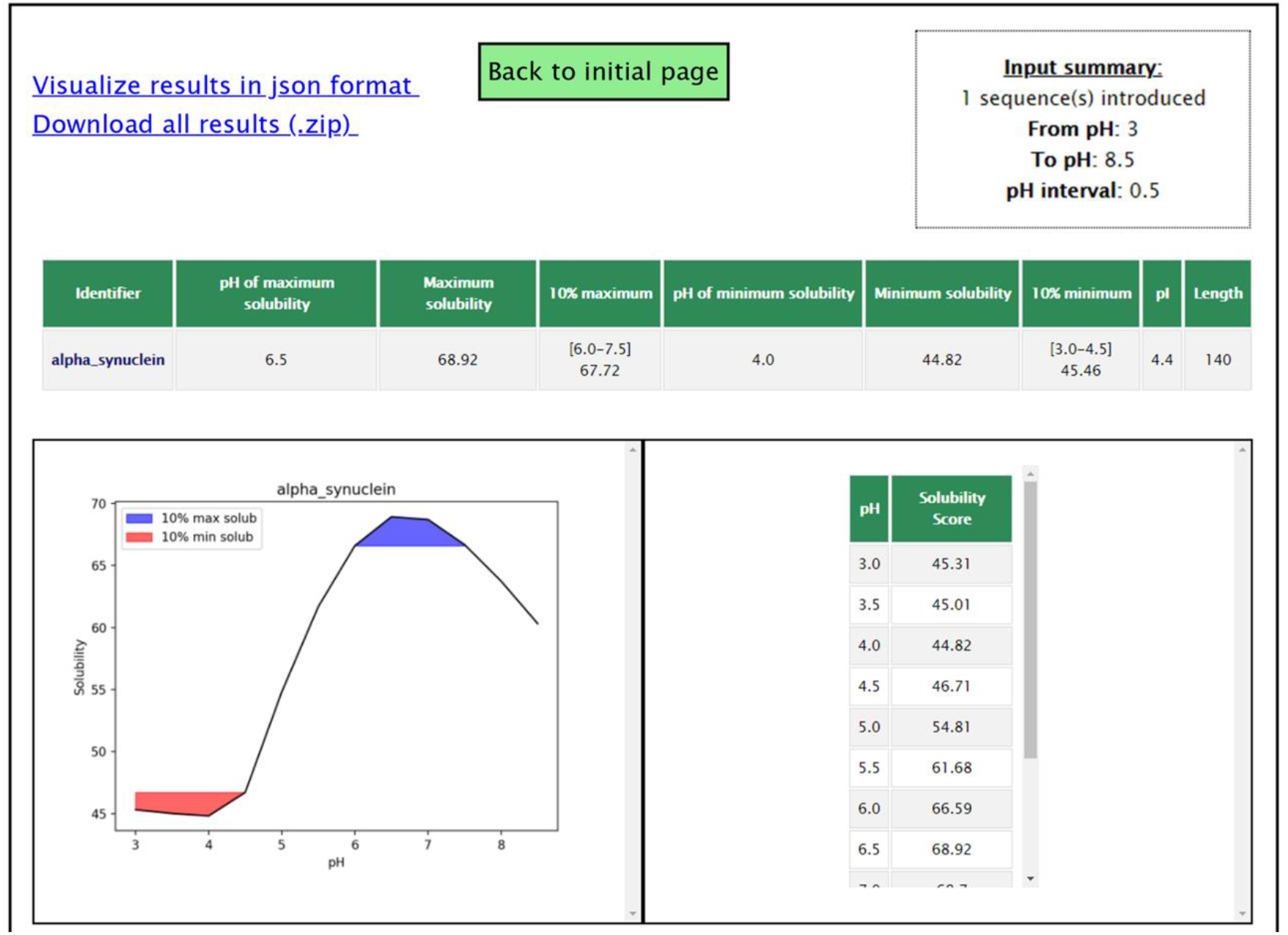
SolupHred output and evaluation: The results page displays two links at the top that allow downloading the numerical results in JSON format or all the project-associated data in a ZIP file. Underneath them, on the left side, a container displays an interactive table with all calculated information for each sequence, showing maximum and minimum solubilities and their respective pHs. On the right, a graphical representation displays solubility variations in the requested pH interval, where the pH of 10% maximum solubility are represented in blue and the pH for the 10% most aggregation-prone in red.

For multiple sequences or a unique pH analysis, the output will be simplified, consisting of a table summarizing the results (Note 9). Either way, all the calculation-derived data can be obtained in a machine-readable way by pressing the upper *“Visualize results in JSON format*” link or downloading a compressed ZIP file with all the generated information for the project.

### 3.2 DispHred web server

#### 3.2.1 Frontpage and running options

DispHred main page shares some common aspects with those described for SolupHred. Its upper edge displays four different sections (Submission, Help, References, and Contact) with a function analogous to those described in Subsection 3.1.1. The main page corresponds to the submission subsection (Figure 5).

**Figure 5.**
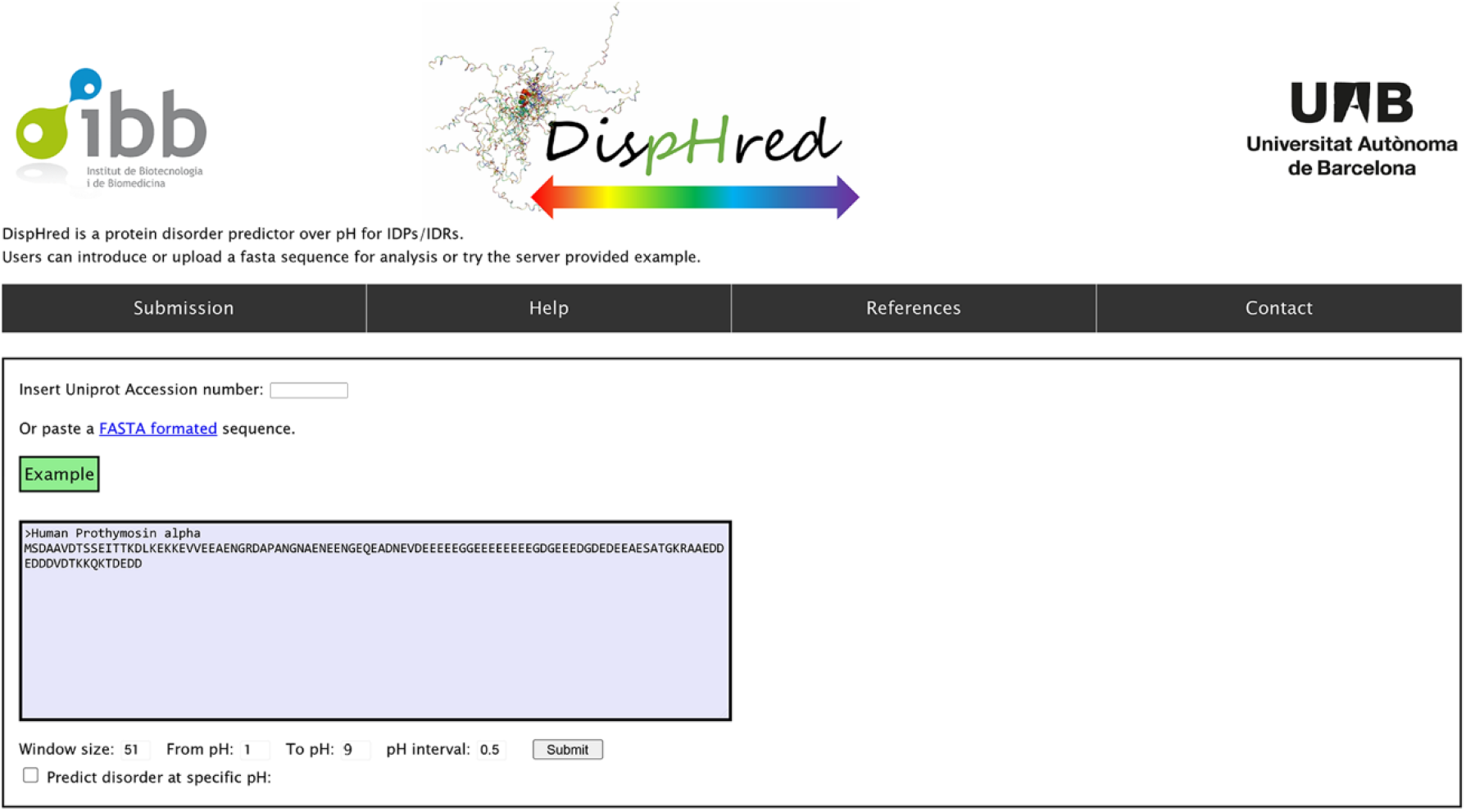
DispHred front page: DispHred displays four clickable headers *Submission, Help, References,* and *Contact.* Under *Submission,* users can paste their FASTA-formatted sequence in the textbox or insert a valid UniprotKB Accession number. DispHred works by default by checking disorder in a range of pHs but allows users to perform predictions at a specific pH. By default, a 51-residue sliding window is populated, but users can personalize its length by changing the value in the textbox. By clicking the *Example* button, the server will pre-populate the human Prothymosin alpha sequence and a pH range for which it undergoes order-disorder transition.

DispHred requires users to input either a FASTA-formatted sequence (Notes 5 and 6) on the provided window or a valid Uniprot Accession number on the textbox (Notes 10 and 11). A 51 amino acid sliding window is selected by default (Note 4), but users can modify it according to their specific needs. The server allows to forecast the pH-dependent disorder in a defined pH interval or, by clicking the *“Predict disorder at a specific pH”,* to limit the analysis at a selected pH. The pH to start and end the analysis and the gradual increment between them can be specified in the text areas *“From pH”, “To pH”* and *“pH interval*” correspondingly. Users can press *Example* to examine the predictions for Human Prothymosin alpha protein in a range of pH that includes those for which the folded-disorder transition occurs [32]. Pressing *Submit* will start the calculations with the precise options specified for the analysis.

#### 3.2.2 Output and evaluation

DispHred redirects users to the results page with the report for the specified analysis. For calculations made through a range of pH, the main output is composed of a table summarizing numerical data, an easy-to-interpret chart, and a color-coded resume of pH-dependent Dis_pH_ scores along the sequence (Figure 6). The table on the left side of the page shows how the Dis_pH_ score, hydrophobicity, and net charge change at the requested pH values. Furthermore, users can click the numerical value for each pH, and a plot with the Dis_pH_ score variation along the sequence will pop up. On the right side of the page, the main chart shows the variation of Dis_pH_ score for the whole sequence along with the range of analyzed pHs. This is accompanied by a dashed red line and two dash-dotted orange lines indicating the order-disorder boundary condition and the confidence interval marked by the SVM. Values above these boundary lines are predicted to be ordered, while those below are predicted to be disordered. This plot allows to interpret the prediction and to recognize at a glance any pH-conditioned order-disorder transition. Below these elements, users can check which protein stretches undergo order-disorder transitions in the specified pH range through a color-coded sequence representation.

**Figure 6.**
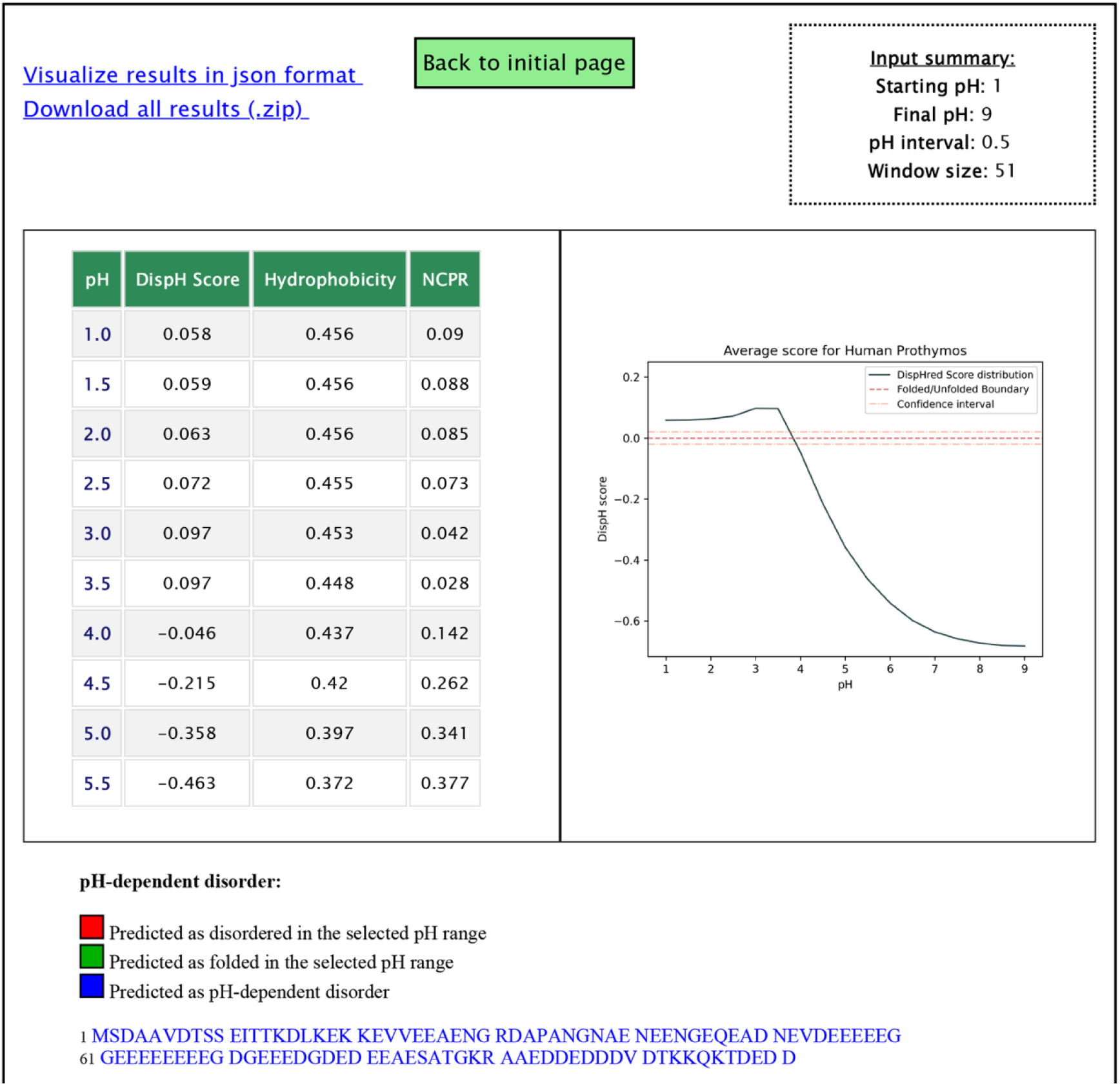
DispHred output and evaluation: In the upper part of the output page, two links are displayed to download the results in JSON format or all the project-generated information in a compressed ZIP file. Below, an interactive table summarizes the results for each pH range, specifying the Dis_pH_ score, hydrophobicity and NCPR. Clicking on each pH identifier will open a per-residue graph for the entire sequence. On its right, a graph represents the Dis_pH_ score variation in the pH interval. A dashed red line and two dash-dotted orange lines indicate the order-disorder boundary condition and the confidence interval. Positive Dis_pH_ scores predict proteins to be ordered, whereas negative Dis_pH_ scores predict them to be disordered. Finally, at the bottom of the results page, a colorimetric scheme shows the pH-dependent disorder in the pH interval accross the sequence.

Alternatively, when analyzing a protein at a specific pH, DispHred will display a first table with the calculated Dis_pH_ score, hydrophobicity, and net charge and a second table showing the calculated Dis_pH_ score along the sequence, accompanied by a plot representing the sequence profile.

Users can retrieve all numerical data in a JSON object or download all the generated output in a compressed ZIP file.

## 4 Concluding remark

The protein environment is a necessary player in conditional disorder and protein aggregation phenomena. For example, environmental pH affects both hydrophobicity and net charge. By modeling how pH impacts these two physicochemical properties, we could anticipate conditional-disorder and self-assembly propensities in IDPs. We next implemented the derived algorithms into the SolupHred and DispHred web servers. These computational approaches were designed to be a fast, cost-effective, and easy to use way to help in further understanding the dynamic nature of IDPs, the mechanisms by which they convert into pathogenic forms, and assist the design of synthetic IDPs, which could transition at specific pHs. They are also expected to help in providing new information on the phenomenon of conditional folding across species or the adaptations of IDPs in organisms living under extreme pH conditions. We expect similar approximations intended to model other environmental conditions being incorporated into state-of-the-art prediction methods in the following years, portraying their predictions into more real-life scenarios.

## 5 Notes

1. SolupHred is trained on IDPs; therefore, the introduced sequences will be assumed to remain disordered in the pH range of analysis. We suggest using DispHred to confirm that the query proteins do not undergo pH-dependent order-disorder transitions on the requested pH range.
2. SolupHred calculates the average lipophilicity of a sliding window and assigns the value to the residue in the center of the window. This window size is defined relative to the protein length: windows of 5 residues are used for proteins shorter than 75 amino acids, 7 for proteins in the 75-174 interval, 9 for 175-299 residues, and, finally, 11 for longer than 300. The resulting values are used to calculate a mean value of global protein lipophilicity.
3. Some proteins’ solubility remains almost constant, showing only modest variations over a range of pHs. SolupHred takes it into account and retrieves the pH ranges with the 10% lowest and highest scores.
4. DispHred uses a 51-residue window by default but allows users to tweak this value according to the scenario that best fits their use case. In instances when the selected window size is larger than the protein length, DispHred will use the complete sequence as a unique window.
5. Both web servers are designed to correct the most common mistakes in submitting FASTA sequences, such as blank spaces or extra line breaks.
6. The server only allows working with the standard 20 proteinogenic amino acids. An error page will be displayed for nonstandard or ambiguous amino acids.
7. SolupHred web server allows pH input to be in the 1 to 14 range, but its performance has not been validated at extreme pH values.
8. SolupHred sets a default step size of 0.5 units of pH. This value can be increased to 2 units of pH, although such high step sizes are not recommended, as any solubility maximum or minimum in between could be missed. On the opposite side, even steps below 0.1 units of pH usually render similar results; SolupHred allows users to perform calculations with ranges down to 0.01 units of pH, if required.
9. SolupHred restricts the automatic generation of graphs up to a maximum of 9 sequences, to increase the velocity of large-scale analysis.
10. If a valid Uniprot Accession number is inserted, the server will retrieve the most updated FASTA entry for the identifier making use of Uniprot’s REST API [29].
11. DispHred is designed to be used with one sequence per analysis. We are currently developing a novel platform to scan large datasets. Meanwhile, we encourage to contact us for these ventures.
12. The graph contains a red line indicating the boundary condition and two orange lines indicating the confidence limits.

## Acknowledgments

This work was funded by the Spanish Ministry of Economy and Competitiveness BIO2016-783-78310-R to S.V, by the Spanish Ministry of Science and Innovation (MICINN) PID2019-105017RB-I00 to S.V. by ICREA, ICREA-Academia 2015 and 2020 to S.V. J. S. was supported by the MICINN (FPU17/01157). C.P.G. was supported by the Secretariat of Universities and Research of the Catalan Government and the European Social Fund (2021 FI_B 00087 to C.P.G.).

